# Glycan-masking spike antigen in NTD and RBD elicits broadly neutralizing antibodies against SARS-CoV-2 variants

**DOI:** 10.1101/2021.11.01.466834

**Authors:** Wei-Shuo Lin, I-Chen Chen, Hui-Chen Chen, Yi-Chien Lee, Suh-Chin Wu

**Affiliations:** Institute of Biotechnology, National Tsing Hua University, Hsinchu 30013, Taiwan; Department of Infectious Diseases, Fu Jen Catholic University Hospital, New Taipei City 242062, Taiwan; Department of Medical Science, National Tsing Hua University, Hsinchu 30013, Taiwan

**Keywords:** SARS-CoV-2, variant, glycan masking, vaccine, COVID-19

## Abstract

Glycan-masking the vaccine antigen by mutating the undesired antigenic sites with an additional N-linked glycosylation motif can refocus B-cell responses to desired/undesired epitopes, without affecting the antigen’s overall-folded structure. This study examine the impact of glycan-masking mutants of the N-terminal domain (NTD) and receptor-binding domain (RBD) of SARS-CoV-2, and found that the antigenic design of the S protein increases the neutralizing antibody titers against the Wuhan-Hu-1 ancestral strain and the recently emerged SARS-CoV-2 variants Alpha (B.1.1.7), Beta (B.1.351), and Delta (B.1.617.2). Our results demonstrated that the use of glycan-masking Ad-S-R158N/Y160T in the NTD elicited a 2.8-fold, 6.5-fold, and 4.6-fold increase in the IC-50 NT titer against the Alpha (B.1.1.7), Beta (B.1.351) and Delta (B.1.617.2) variants, respectively. Glycan-masking of Ad-S-D428N in the RBD resulted in a 3.0-fold and 2.0-fold increase in the IC50 neutralization titer against the Alpha (B.1.1.7) and Beta (B.1.351) variants, respectively. The use of glycan-masking in Ad-S-R158N/Y160T and Ad-S-D428N antigen design may help develop universal COVID-19 vaccines against current and future emerging SARS-CoV-2 variants.

## Introduction

Severe acute respiratory syndrome coronavirus 2 (SARS-CoV-2) is an RNA virus that has a higher mutation rate than host cells or DNA viruses, because of its inherent ability to generate a quasi-species mutant genome of variant viruses during replication (1). Multiple SARS-CoV-2 variants have already emerged and are circulating globally, four of which are a cause of serious public health concern, also known as variants of concern: (i) Alpha (B.1.1.7 lineage) variant which originated in the United Kingdom (UK), (ii) Beta (B.1.351 lineage) variant which originated in South Africa, (iii) Gamma (P.1 lineage) variant which originated in Brazil, and (iv) Delta (B.1.617.2 lineage) variant which originated in India (2). The Alpha (B.1.1.7) variant encodes an S protein with nine mutations (del 69-70, Del 144, N501Y, A570D, D614G, P681H, T716I, S982A, and D1118H), of which N501Y is in the receptor-binding domain (RBD). The Beta (B.1.351) variant encodes an S protein with nine mutations (L18F, D80A, D215G, Del 241-243, K417N, E484K, N501Y, D614G and A701V, three of which (K417N, E484K and N501Y) are in the RBD. The Gamma P.1 variant encodes an S protein with 12 mutations (L18F, T20N, P26S, D138Y, R190S, K417N/T, E484K, N501Y, D614G, H655Y, T1027I, and V1176F), two of which are in the RBD (E484K, and N501Y). The Delta (B.1.617.2) variant encodes an S protein with ten mutations (T19R, G142D, del 156, del 157, R158G, L452R, T478K, D614G, P681R, and D950N), two of which are in the RBD (L452R and T478K). The immunity-evading mutations in the Beta (B1.351) variant include E484K in the RBD of the S protein (3), while those in the Delta (B.167.2) variant include L19R, del 157, del 158, L452R, and T478K in the S protein (4-6). In particular, the RBD mutations K417, L452, E484 and N501 may directly form a new interaction with the human angiotensin-converting enzyme 2 (ACE2) receptor (7-8). Thus, the emerging SARS-CoV-2 B.1.351 variant can evade antibody-mediated immunity without affecting the virus fitness and disease, as recently reported using convalescent plasma, vaccine-induced sera, and monoclonal antibodies (9-15). Whether the current authorized or approved vaccines can still provide effective protection against current and future emerging SARS-CoV-2 variants remains unclear.

Glycan-masking the vaccine antigen by mutating the undesired antigenic sites with an additional N-linked glycosylation motif can refocus the B-cell responses to the desired/undesired epitopes, without affecting the antigen’s overall-folded structure (16-17). This antigen design strategy has been used to develop vaccines against human immunodeficiency virus (HIV)-1 (18-23), influenza virus (24-27), dengue and Zika viruses (28), and Middle East respiratory syndrome coronavirus (29). In this study, we used an adenovirus (Ad) vector encoding the full-length *S* gene of the SARS-CoV-2 Wuhan-Hu-1 isolate with a series of site-specific glycan-masking mutations on the N-terminal domain (NTD) and RBD in a mouse immunization model, and then investigated the breadth of neutralizing antibodies elicited against SARS-CoV-2 and its Alpha (B.1.1.7), Beta (B.1.351), and Delta (B.1.617.2) variants. These results can provide useful information for the further development of universal coronavirus disease 2019 (COVID-19) vaccines against current and future emerging SARS-CoV-2 variants.

## Materials and Methods

### Cell lines

HEK293A and HEK293T cells were obtained from the Bioresource Collection and Research Center (BCRC), Taiwan. These cells were grown in Dulbecco’s modified Eagle medium (DMEM) (Thermo Scienific) supplemented with 10% fetal bovine serum (FBS) (Gibco) and 100 units/ml penicillin/streptomycin (P/S), and maintained in an incubator at 37°C with 5% CO_2_.

### Selection of glycan-masking sites from the SARS-CoV-2 S protein structure

Selection of the glycan-masking sites was based on the 3D-structure of SARS-CoV-2 S protein (PDB ID:7C2L). The exposed loops or the protruding sites of the exposed loops on the NTD and RBD of the S protein were examined using PyMol (The PyMol Molecular Graphics System, version 4.0; Schrödinger, LLC). Glycan-masking sites that are less than 5 Å from the native glycans and RBD were discarded.

### Preparation of Ad vectors expressing SARS-CoV-2 S gene and glycan-masking mutants

The human codon-optimized S gene of SARS-CoV2 (Wuhan-Hu-1 isolate, accession number MN908947.3) was obtained from GenScript. Site-directed mutagenesis was used to produce the glycan-masking S mutant genes, with the addition of an N-linked glycosylation motif at the S protein residues 135N/N137T, R158N/Y160T, N354/K356T, N370/A372T, G413N, D428N and H519N/P521T. Wild-type S and glycan-masking S genes were first cloned into the pENTR1A vector (Invitrogen), and then cloned into the adenoviral plasmid pAd/CMV/V5-DEST (Invitrogen) using LR ClonaseTM II Enzyme Mix (Invitrogen) to produce the Ad plasmid expressing SARS-CoV-2 S gene. To obtain Ad particles, the Ad plasmids were cleaved with Pac I restriction enzyme to expose the inverted terminal repeats and then transfected into 293A cells separately using TurboFect transfection reagent (Fermentas). After 10-15 d, once the cytopathic effects were visible, the transfected cells and culture media were collected. The cells were disrupted by means of three freeze-thaw cycles to release the intracellular viral particles, and the supernatants of the cell lysates were collected by centrifugation (3000 rpm, 15 min, 4°C) to obtain the Ad vectors expressing the SARS-Co-V-2 S proteins. To prepare higher titers, the virus was concentrated using a 30-kDa Amicon Ultra-15 Centrifugal Filter (Millipore). The viral stocks were stored at -80°C. To determine the Ad titers, HEK293A cells were seeded into 6-well plates at a density of 10^6^ cells/well and incubated at 37°C overnight. The 10-fold serially diluted Ad stocks were then added to each well at 37°C for 24 h. Next, the media containing the diluted Ad vectors were removed, and 3 mL/well of DMEM containing 0.4% agarose and 100 U/ml P/S was added to the 6-well plates. The plaques were visibly quantified 7-10 d after the cells were infected with Ad vectors, and the pfu count was noted.

### SDS-PAGE and western blot

HEK293A cells were infected with Ad-S, Ad-S-F135N/N137T, Ad-S-R158N/Y160T, Ad-N354/K356T, Ad-S-N370/A372T, Ad-S-G413N, Ad-S-D428N and Ad-S-H519N/P521T at an MOI = 5 for 48 h, then lysed with Glo Lysis buffer (Promega), and subjected to centrifugation at 12000 × *g* for 5 min at 4°C to remove the cell debris. The lysates were heated at 95°C for 5 min in reducing sample buffer, treated or not treated with PNGase F (BioLabs) for 2 h at 37°C, and resolved using 10% or 7% SDS-PAGE gels. For western blots, nitrocellulose membranes (Millipore) were blocked in 5% (w/v) skim milk for 1 h at room temperature, followed by three washes with Tris-buffered saline containing 0.05% Tween 20 (TBST). SARS-CoV-2 S proteins were probed with anti-SARS-CoV-2 primary antibodies (GTX135356, GeneTex) overnight at 4°C, and detected with horseradish peroxidase (HRP)-conjugated goat anti-rabbit IgG (KPL) for 1 h at room temperature. HRP-catalyzed enhanced chemiluminescence (Millipore) was captured using an X-ray film.

### Mouse immunization

Groups of female BALB/c mice (6 to 8 weeks old) (n=5 per group) were obtained from the National Laboratory Animal Center, Taipei, Taiwan. Groups of female BALB/c mice (6 to 8 weeks old) (n=5 per group) were immunized with Ad-S, Ad-S-F135N/N137T, Ad-S-R158N/Y160T, Ad-S-N370/A372T, or Ad-S-H519N/P521T vectors at 5 × 10^7^ plaque-forming unit (pfu) per dose in PBS (pH 7.4) in the first set of immunization experiments, and immunized with Ad-S, Ad-S-N354/K356T, Ad-S-G413N, and Ad-S-D428N vectors at 1 × 10^8^ pfu per dose in the second set of immunization experiments. Intramuscular injections were administered at weeks 0 and 3. Sera were collected 2 weeks after the second immunization dose.

### Enzyme-linked immunosorbent assay (ELISA)

To measure the SARS-CoV-2 specific total IgG titer in the antisera, recombinant S (Wuhan-Hu-1, catalog number 40589-V08H4), RBD (Wuhan-Hu-1, cat number 40592-V08H), S1 (B.1.1.7 variant, catalog number 40591-VH12), RBD (B.1.1.7 variant, catalog number 40592-V08H82), S1 (B.1.351 variant, cat number 40591-V08H10), RBD (B.1.351 variant, catalog number 40592-V08H85), S1 (B.1.617.2 variant, catalog number 40591-V49H2-B), RBD (B.1.617.2 variant, catalog number 40592-V08H90) proteins were obtained from Sino Biological Inc., and allowed to coat 96-well plates at a concentration of 2μg/mL in coating buffer (100μL/well) overnight at 4°C. Coating buffers were aspirated and washed three times with PBS containing 0.05% Tween 20 (PBST). Each well was blocked with 200μL blocking buffer (1% BSA in PBST) at room temperature for 2 h. Heat-inactivated serum samples were pre-diluted 1:1000, followed by 2-fold serial dilution in dilution buffer (0.05% tween 20 +1% BSA in PBST). The plates were washed three times with 300 μL PBST (PBS with 0.05% Tween-20), and then blocked with 200 μL PBS buffer plus 1% BSA for 2 h at room temperature, followed by three additional washes with 300 μL PBST. Following this, the plates were incubated with 100 μl of HRP) conjugated anti-mouse IgG antibody (1:30000 in dilution buffer) for 1 h at room temperature. After three additional washes with 300 μL PBST, 100 μL of TMB substrate (BioLegend) was added to each well and incubated in the dark for 15 min. The reaction was stopped by the addition of 100 μL of 2 N H_2_SO_4_. The optical density at 450 nm was measured using a TECAN spectrophotometer. The end-point titration values were calculated in terms of a final serial dilution higher than 0.2 optical density value.

### SARS-CoV-2 pseudotyped lentivirus neutralization assay

To produce SARS-CoV-2 pseudoviruses, a plasmid expressing the full-length S protein (Wuhan-Hu-1, B.1.1.7, or B.1.351) of SARS-CoV-2 was co-transfected into HEK293T cells with packaging and reporter plasmids pCMVΔ8.91 and pLAS2w.FLuc.Ppuro (RNAi Core, Academia Sinica), using TransIT-LT1 transfection reagent (Mirus Bio). The medium was harvested and concentrated at 48 h post-transfection, followed by estimation of the pseudovirus titer in terms of the luciferase activity of SARS-CoV2-Spp transduction. Serum samples were serially diluted and incubated with 1,000 TU of SARS-CoV-2-pseudotyped lentivirus in DMEM (supplemented with 1% FBS and 100 U/mL P/S) for 1 h at 37°C. The mixture was then inoculated with an equal volume of 10,000 HEK-293T cells stably expressing the ACE2 gene in 96-well plates, which were seeded one day before infection. The culture medium was replaced with fresh complete DMEM (supplemented with 10% FBS, 100 U/mL P/S) at 16 h post-infection and the cells were then continuously cultured for another 48 h before being subjected to a luciferase assay (Promega Bright-GloTM Luciferase Assay System). The percentage of inhibition was calculated as the ratio of the loss of luciferase readout (RLU) in the presence of serum to that of the no serum control. The formula used for the calculation was (RLU Control - RLU Serum) / RLU Control. Neutralization titers (IC50) were measured as the reciprocal of the serum dilution required to obtain a 50% reduction in RLU compared to a control containing the SARS-CoV-2 S-pseudotyped lentivirus only. Neutralization curves and IC50 values were analyzed using the GraphPad Prism 5 Software.

### Statistical analyses

Statistical tests for multiple comparison were performed for all groups (except for the PBS control) in case of the ELISA data. The results were analyzed using the nonparametric Kruskal-Wallis test, with corrected Dunn’s multiple comparison test, using GraphPad Prism v6.01. Statistical significance has been expressed as follows: *p < 0.05; **p < 0.01; and ***p < 0.001. Neutralization curves were fitted based on the equation of nonlinear regression log (inhibitor) vs. normalized response -- variable slope using GraphPad Prism v6.01. The IC50 values of the neutralization were obtained from the fitting curves using GraphPad Prism v6.01

## Results

### Design of engineered glycan-masking S antigens in the NTD and RBD for Ad vector immunization

The S protein of SARS-CoV-2 is trimeric, and each monomer comprises of S1 and S2 subunits (30-32). The S1 subunit contains NTD and RBD. To design glycan-masking S antigen(s) for immunization, we used an Ad vector encoding the full-length S gene of the SARS-CoV-2 Wuhan-Hu isolate, by introducing a series of N-linked glycosylation motifs into the S1 region of the S protein, to refocus the antibody responses to the RBD (**Fig. 1A**). The sites of glycan-masking were introduced not only in the RBD, but also in the NTD, as RBD and NTD may spatially interact with each other in the quaternary structure of the intact trimeric S protein (**Fig. 1B**). The exposed loops or the protruding sites of the exposed loops on the NTD and RBD of the 3-D S protein structure (PDB ID: 7C2L) were chosen for the addition on the glycan-masking sites. Seven glycan-masking N-glycan sites were engineered in the Ad-S vector: (#1) Ad-S-F135N/N137T, (#2) Ad-S-R158N/Y160T, (#3) Ad-S-N370/A372T, (#4) Ad-S-H519N/P521T, (#5) Ad-S-N354/K356T, (#6) Ad-S-G413N, and (#7) Ad-S-D428N) (**Fig. 1B**). To characterize the glycan-masking mutations on the S protein, the lysates of HEK293A cells infected with each Ad-S vector were analyzed using 8% SDS-PAGE gels, followed by western blotting with an S1-specific polyclonal antibody. The results indicated the presence of S and S1 in the cell lysates of HEK293A cells infected with Ad-S, Ad-S-F135N/N137T, Ad-S-R158N/Y160T, Ad-S-N354/K356T, Ad-S-N370/A372T, Ad-S-G413N, Ad-S-D428N, and Ad-S-H519N/P521T (**Fig. 2**).

**Figure 1.**
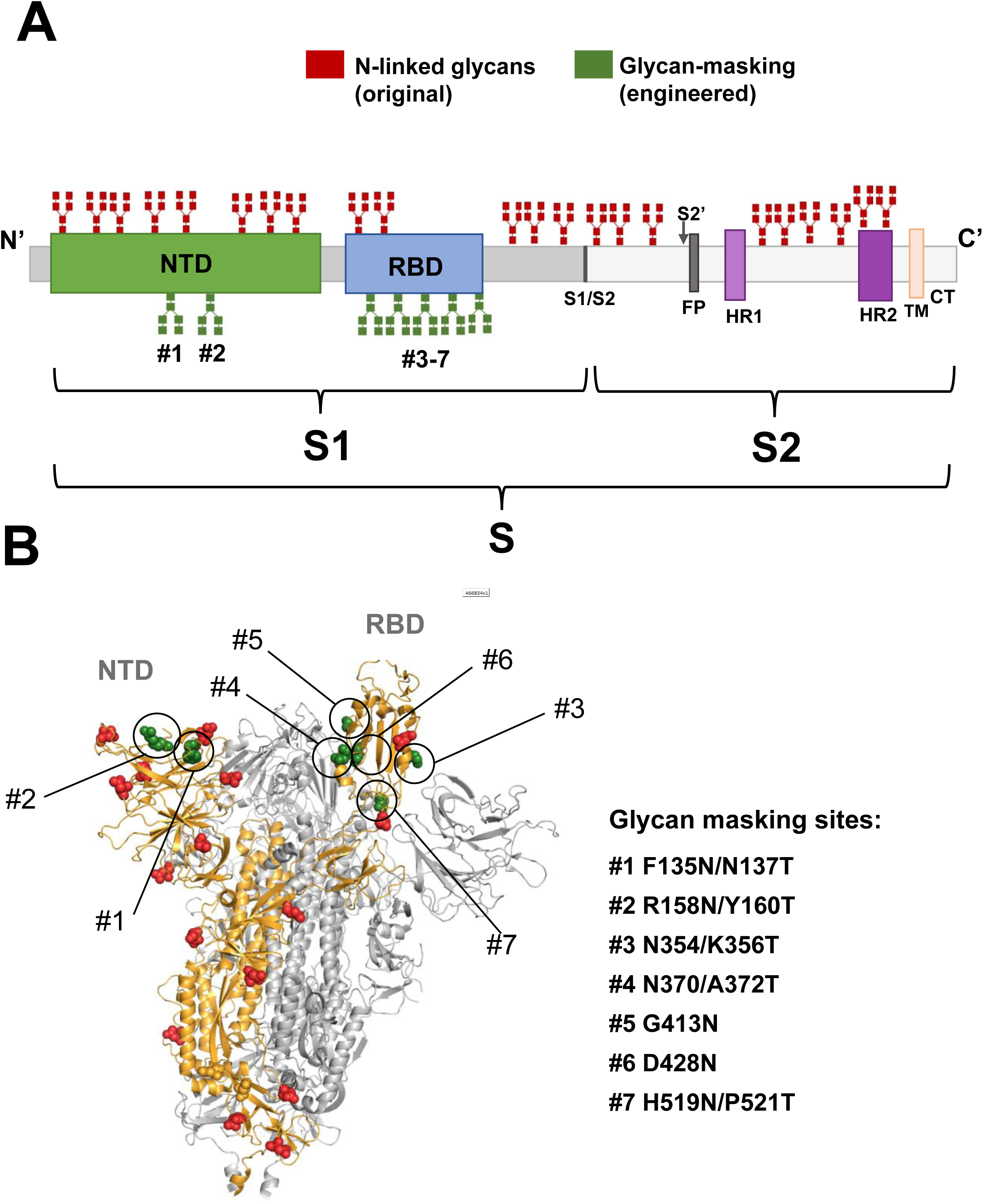
Schematic representation of SARS COV2 spike glycoprotein. **(A)** Positions of the original N-linked glycan (red) and the engineered N-linked glycan (green) amino-acid sequences shown as branches. Domains of the full-length S protein: N-terminal domain (NTD), receptor binding domain (RBD), furin cleavage site (S1/S2), fusion peptide (FP), heptad repeat 1(HR1), heptad repeat 2 (HR2), transmembrane domain (TM), and cytoplasmic tail (CT); **(B)** The seven engineered N-linked glycan sites of (#1) F135N/N137T, (#2) R158N/Y160T, (#3) N354/K356T, (#4) N370/A372T, (#5) G413N, (#6) D428N, and (#7) H519N/P521T are shown in the intact trimeric S structure.

**Figure 2.**
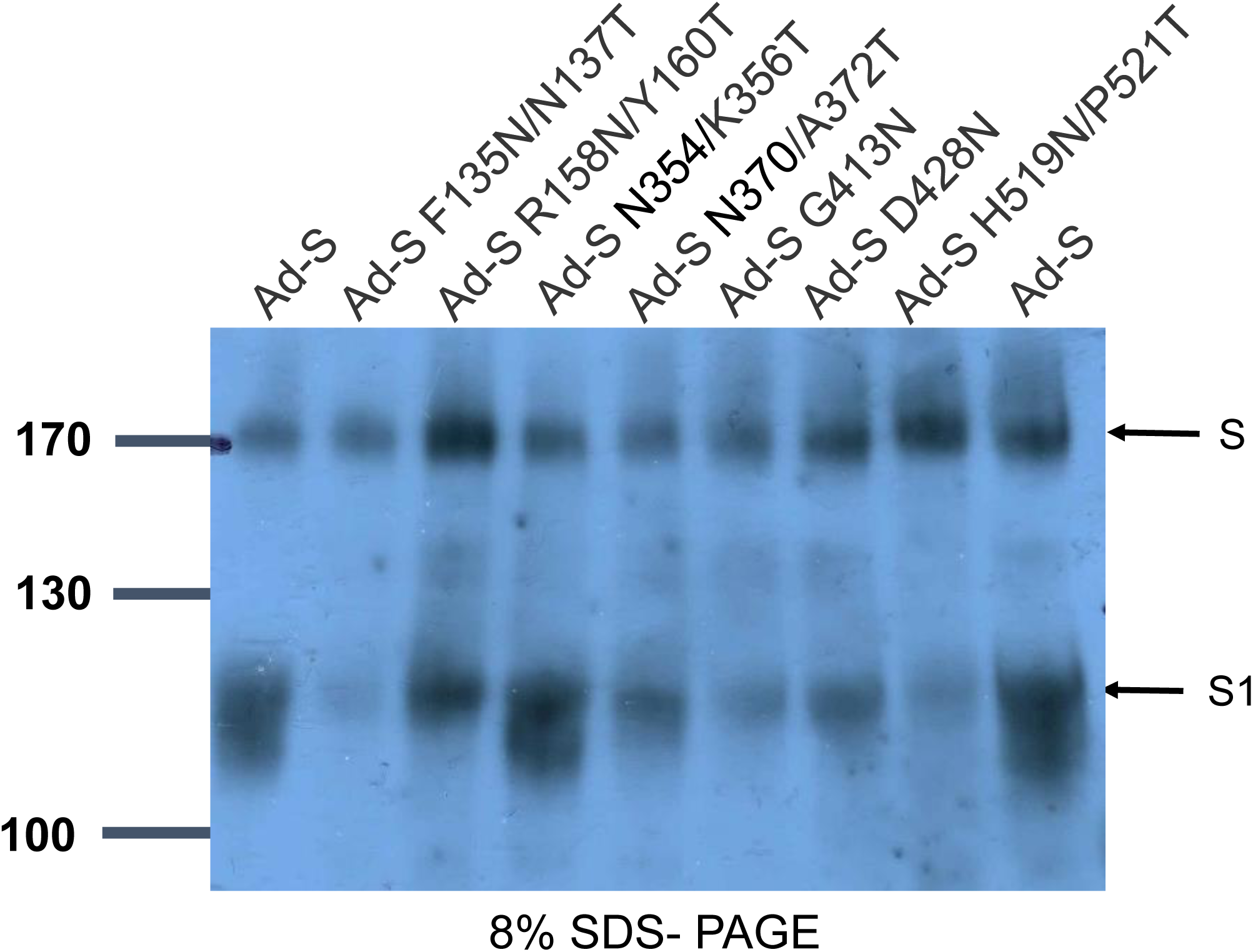
Characterization of Ad-S glycan-masking mutants using SDS-PAGE and western blotting. Cell lysates of HEK-293A cells infected with Ad-S, Ad-S-F135N/N137T, Ad-S-R158N/Y160T, Ad-S-N354/K356T, Ad-S-N370/A372T, Ad-S-G413N, Ad-S-D428N, and Ad-S-H519N/P521T and analyzed using 8% SDS-PAGE and western blotting with an S1-specific polyclonal antibody. The results indicated the presence of S and S1 in the cell lysates.

### Anti-S, anti-RBD, and pseudo-neutralizing antibody titers against the Wuhan-Hu-1 ancestral strain

To investigate the antibody responses elicited by these Ad-S vectors, groups of BALB/c mice (n=5 per group) were intramuscularly immunized with two doses of each Ad vector at 5 × 10^7^ pfu per dose for the first set of immunization experiments and at 1 × 10^8^ pfu per dose for the second set of immunization experiments all in a 3-week interval; PBS-immunized animals were used as control (**Fig. 3A**). Sera from these immunized groups were collected after 2 weeks of the second dose and analyzed for the levels of anti-S, anti-RBD, and neutralizing antibodies against the original SARS-CoV-2 Wuhan-Hu-1 isolate. Two separate sets of immunization experiments were conducted in this study: (i) with Ad-S, Ad-S-F135N/N137T, Ad-S-R158N/Y160T, Ad-S-N370/A372T, and Ad-S-H519N/P521T, and (ii) with Ad-S, Ad-S-N354/K356T, Ad-S-G413N, and Ad-S-D428N. In the first set of immunization experiments, the results indicated that the anti-S IgG titer elicited in the Ad-S-F135N/N137T-immunized group was significantly lower than those elicited in the wild-type Ad-S and Ad-S-R158N/Y160T-immunied groups (**Fig. 3B**). A reduced (but not statistically significant) titer of anti-RBD IgG antibodies was also observed for the Ad-S-F135N/N137T-immunized group, as compared to the other four immunized groups (**Fig. 3C**). Pseudo-neutralizing antibody titers from the pooled sera of each group (n =5 mice per group) were determined using the SARS-CoV-2 S (Wuhan-Hu-1)-pseudotyped lentivirus assay in triplicate. In the first set of immunization experiments, Ad-S, Ad-S-F135N/N137T, Ad-S-R158N/Y160T, Ad-S-N370/A372T, and Ad-S-H519N/P521T-immunized groups showed dose-response neutralization, while the PBS-immunized control group did not (**Fig. 3D)**. Antisera from the Ad-S-R158N/Y160T-immunized group showed increased neutralization potency, as compared to those of the Ad-S and Ad-S-N370/A372T-immunized groups, and the lower levels of the Ad-S-F135N/N137T and Ad-S-H519N/P521T-immunized groups (**Fig. 3D**). The corresponding IC-50 titer elicited in the Ad-S-R158N/Y160T-immunied group against the Wuhan-Hu-1 ancestral strain was approximately 2.4-fold higher than that elicited in the wild-type Ad-S-immunized group (**Fig. 3E**). No significant differences were observed in the anti-S and anti-RBD titers in the second set of immunization experiments with the Ad-S, Ad-S-N354/K356T, Ad-S-G413N, and Ad-S-D428N-immunized groups **(Fig. 3E-F)**. Dose-dependent pseudo-neutralization curves were observed for the Ad-S, Ad-S-N370/K356T, Ad-S-G413N, and Ad-S-D428N-immunized groups, but not for the PBS-immunized control **(Fig. 3G)**. The IC-50 titers of the Ad-S-N354/K356T and Ad-S-D428N-immunized groups against the Wuhan-Hu-1 ancestral strain were approximately 2.5- and 2.8-fold higher, as compared to that of the wild-type Ad-S-immunized group (**Fig. 3H**). These results indicated that the glycan-masking Ad-S-R158N/Y160T in NTD and glycan-masking Ad-S-N354/K356T and Ad-S-D428N in RBD elicited a 2.4-, 2.5-, and 2.8-fold increase, respectively, in the pseudo-neutralization IC50 titer against the Wuhan-Hu-1 ancestral strain.

**Figure 3.**
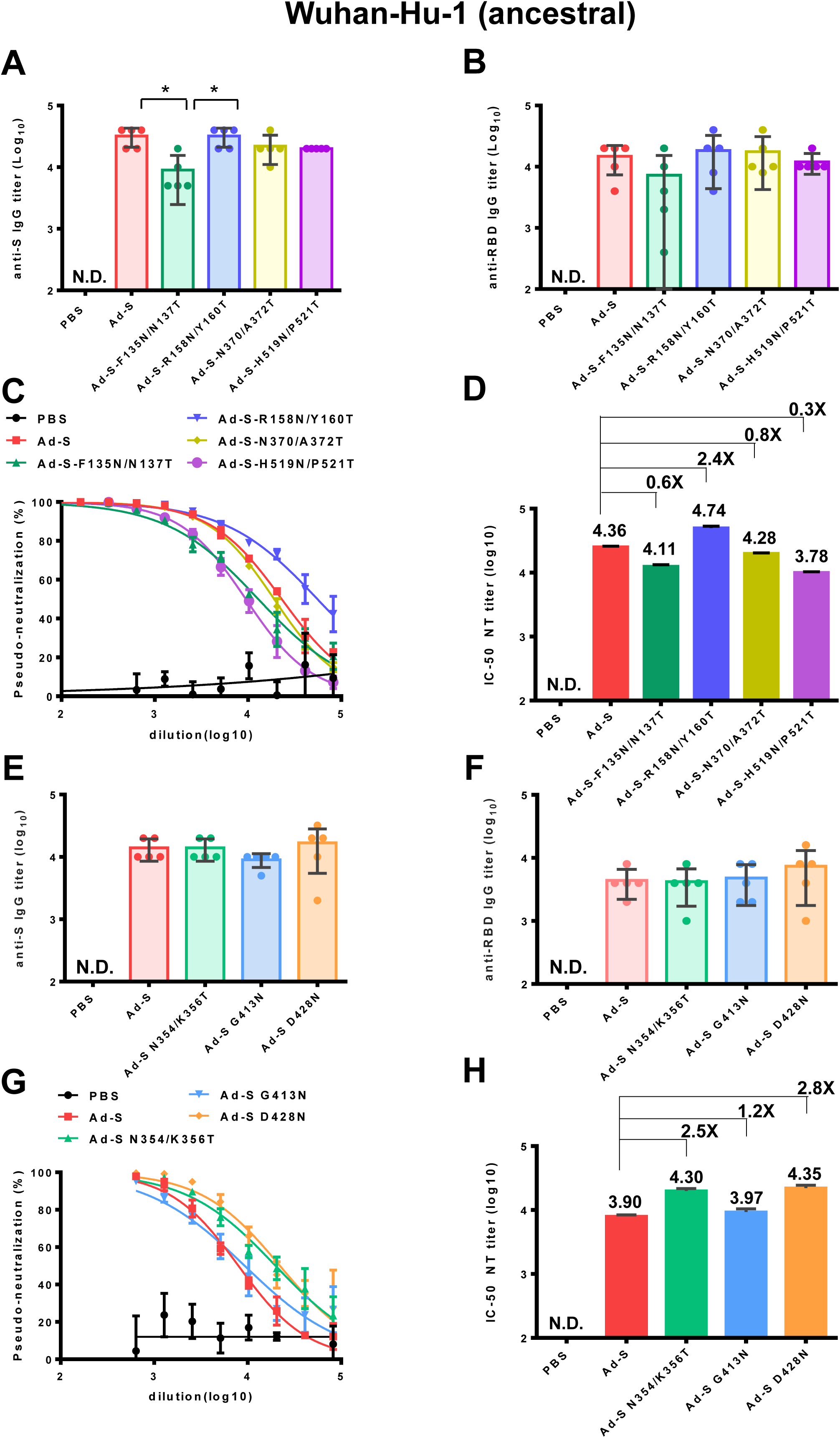
Antibody responses elicited by Ad-S glycan-masking mutants against the SARS-CoV-2 Wuhan-Hu-1 strain. Ad-S immunization regimens in the first and second sets of immunization experiments: (**A)** antisera for anti-S IgG titers from each group of mice (n=5) and tested individually in the first set of immunization experiments; **(B)** antisera for anti-RBD IgG titers from each group of mice (n=5) and tested individually in the first set of immunization experiments; **(C)** the dose-response pseudo-neutralization curves for the pooled sera of each immunized group of mice (n=5) and measured in triplicate the first set of immunization experiments; **(D)** IC50 NT titer for neutralizing antibodies against the Wuhan-Hu-1 ancestral strain in the first set of immunization experiments; (**E)** antisera for anti-S IgG titers from each group of mice (n=5) and tested individually in the second set of immunization experiments; **(F)** antisera for anti-RBD IgG titers from each group of mice (n=5) and tested individually in the second set of immunization experiments; **(G)** the dose-response pseudo-neutralization curves for the pooled sera of each immunized group of mice (n=5) and measured in triplicate the second set of immunization experiments; **(H)** IC50 NT titer for neutralizing antibodies against the Wuhan-Hu-1 ancestral strain in the second set of immunization experiments. Fold-changes of the Ad-S glycan-masking mutant IC-50 NT titers against the wild-type Ad-S titer on a linear scale indicated. Statistical tests for multiple comparison of anti-S and anti-RBD IgG titers among the Ad-S immunization groups were performed using the nonparametric test with Kruskal-Wallis with corrected Dunn’s multiple comparison. The statistical significance is expressed as follows: *p < 0.05. Neutralization curves were fitted based on the equation of nonlinear regression log (inhibitor) vs. normalized response -- variable slope using GraphPad Prism v6.01. The IC50 values of neutralization were obtained from the fitting curves using GraphPad Prism v6.01. Error bars are plotted as standard deviation from the mean value. Not detectable for N.D.

### Anti-S1, anti-RBD, and pseudo-neutralizing antibody titers against SARS-CoV-2 Alpha (B.1.1.7), Beta (B.1.351), and Delta (B.1.617.2) variants

To further study the neutralization against SARS-CoV-2 variants, the titers of anti-S1, and anti-RBD IgG, and neutralizing antibodies against the Alpha (B.1.1.7), Beta (B.1.351), and Delta (B.1.617.2) variants were measured using the same antisera. In the first set of immunization experiments in the Ad-S, Ad-S-F135N/N137T, Ad-S-R158N/Y160T, Ad-S-N370/A372T, and Ad-S-H519N/P521T-immunized groups, we found that the anti-S1 IgG titers against Alpha (B.1.1.7) variant in the Ad-S-F135N/N137T-immunized group were lower than the wild type Ad-S and Ad-S-N370/A372T-immunized groups (**Fig. 4A**). A reduced (but not statistically significant) titer of anti-RBD IgG antibodies against Alpha (B.1.1.7) variant was observed for the Ad-S-F135N/N137T-immunized group, as compared to the other four immunized groups (**Fig. 4B**). The pseudovirus neutralization curve of the Ad-S-R158N/Y160T-immunized group against the Beta (B.1.1.7) variant was more potent than those of the wild-type Ad-S-immunized group and the three other immunization groups (**Fig. 4C**), with an approximately 2.8-fold increase in the neutralization IC50 titer, as compared to that of the wild-type Ad-S-immunized group (**Fig. 4D**). In the second set of immunization experiments in the Ad-S, Ad-S-N354/K356T, Ad-S-G413N, and Ad-S-D428N-immunized groups, no significant differences were observed in the anti-S1 and anti-RBD titers among these four Ad immunization groups **(Fig. 4E-F)**. However, the Ad-S-D428N-immunized group displayed more potent neutralization against the Beta (B.1.351) variant than the three other groups (**Fig. 4G**), resulting in a 3.0-fold increase in the neutralization IC50 titer, as compared to that of the wild-type Ad-S-immunized group (**Fig. 4H**). Therefore, the glycan-masking Ad-S-R158N/Y160T in NTD and glycan-masking Ad-S-D428N in RBD were found to elicit increased titers of neutralizing antibodies against the Alpha (B.1.1.7) variant.

**Figure 4.**
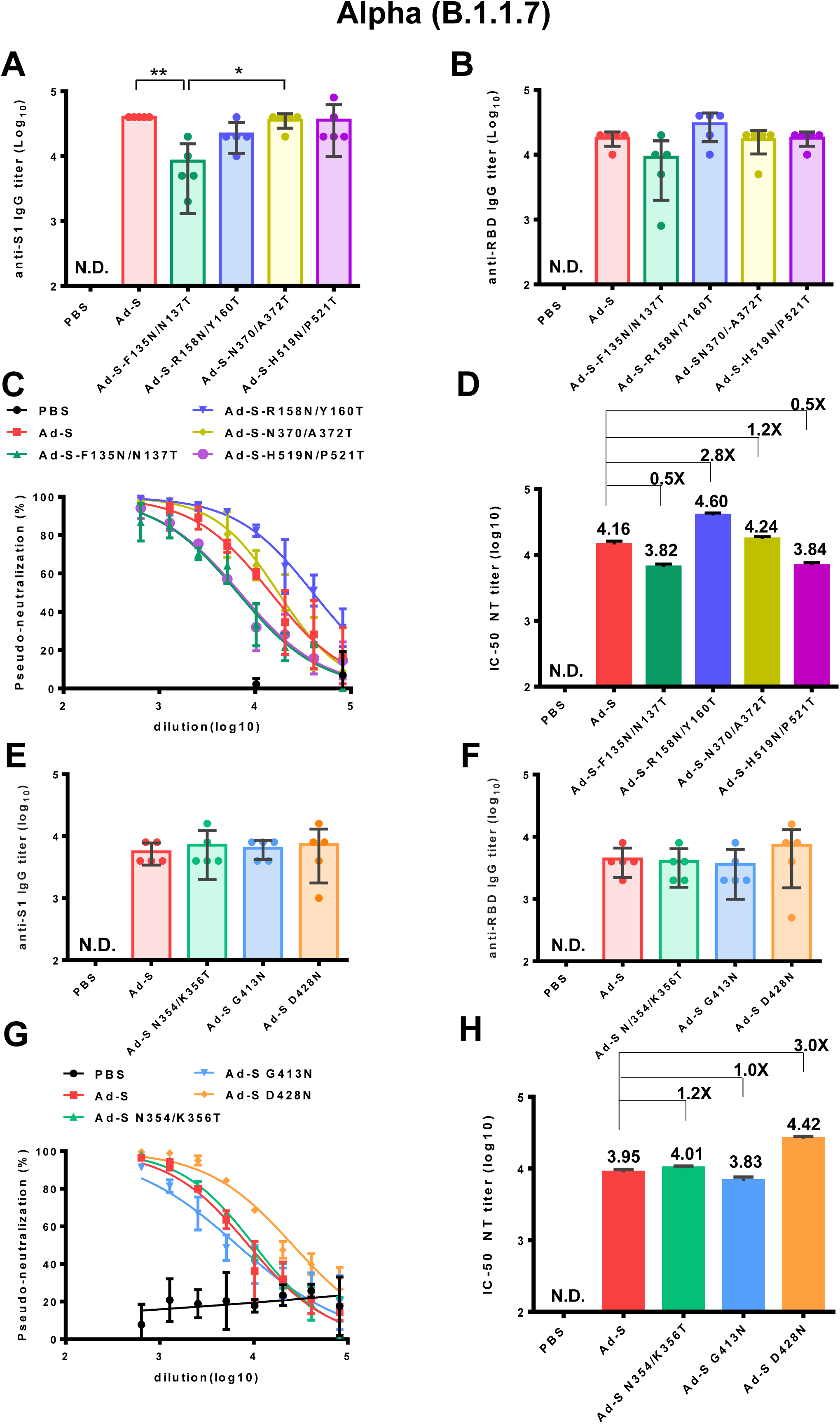
Antibody responses elicited by Ad-S glycan-masking mutants against SARS-CoV-2 Alpha (B.1.1.7) variant. **(A)** antisera for anti-S1 IgG titers from each group of mice (n=5) and tested individually in the first set of immunization experiments; **(B)** antisera for anti-RBD IgG titers from each group of mice (n=5) and tested individually in the first set of immunization experiments; **(C)** the dose-response pseudo-neutralization curves for the pooled sera of each immunized group of mice (n=5) and measured in triplicate in the first set of immunization experiments; **(D)** IC-50 NT titer for neutralizing antibodies against Alpha (B.1.1.7) variant; **(E)** antisera for anti-S1 IgG titers from each group of mice (n=5) and tested individually in the second set of immunization experiments; **(F)** antisera for anti-RBD IgG titers from each group of mice (n=5) and tested individually in the second set of immunization experiments; **(G)** the dose-response pseudo-neutralization curves for the pooled sera of each immunized group of mice (n=5) and measured in triplicate for the second set of immunization experiments; **(H)** IC-50 NT titer for neutralizing antibodies against Alpha (B.1.1.7) variant for the second set of immunization experiments. Fold-changes of the Ad-S glycan-masking mutant IC-50 NT titers against the wild-type Ad-S (Alpha, B.1.1.7 variant) titer on a linear scale are indicated. Statistical tests for multiple comparison of anti-S1 and anti-RBD titers among Ad-S immunization groups were performed using the nonparametric test with Kruskal-Wallis with corrected Dunn’s multiple comparison. The statistical significance is expressed as follows: *p < 0.05 and **p < 0.01. Neutralization curves were fitted based on the equation of nonlinear regression log (inhibitor) vs. normalized response -- variable slope using GraphPad Prism v6.01. The IC50 values of neutralization were obtained from the fitting curves using GraphPad Prism v6.01. Not detectable for N.D. Error bars are plotted as standard deviation from the mean value.

In the case of the Beta (B.1.351) variant in the first set of immunization experiments in the Ad-S, Ad-S-F135N/N137T, Ad-S-R158N/Y160T, Ad-S-N370/A372T, and Ad-S-H519N/P521T-immunizd groups, the Ad-S-F135N/N137T-immunized group had a significantly lower IgG titer of anti-S1 antibodies than those in the wild type Ad-S and Ad-S-N370/A372T-immunized groups (**Fig. 5A**). The anti-RBD IgG titer in the Ad-S-F135N/N137T-immunized group was lower, as compared to the titers in the Ad-S-R158N/Y160T and Ad-S-H519N/P521T-immunizd groups (**Fig. 5B**). Both the Ad-S-R158N/Y160T and Ad-S-N370/A372T groups showed increased potency for pseudo-neutralization against the Beta (B.1.351) variant, as compared to the three other groups (Ad-S, Ad-S-F135N/N137T, and Ad-S-H519N/P521T-immunized) (**Fig. 5C**); the neutralization IC-50 titer was approximately 6.5-fold and 2.8-fold higher in the glycan-masking Ad-S-R158N/Y160T and Ad-S-N370/A372T-immunized groups, respectively (**Fig. 5D**). In the second set of immunization experiments in the Ad-S, Ad-S-N354/K356T, Ad-S-G413N, and Ad-S-D428N-immunized groups, no significant differences were observed in the anti-S1 and anti-RBD titers among these four Ad immunization groups **(Fig. 5E-F)**. The Ad-S-D428N-immunized group was more potent in neutralizing the Beta (B.1.351) variant than the other three groups (**Fig. 5G**), resulting in a 2.0-fold increase in the neutralization IC50 titer, as compared to that of the wild-type Ad-S-immunized group (**Fig. 5H**). Therefore, immunization with the glycan-masking Ad-S-R158N/Y160T in NTD and glycan-masking Ad-S-N370/A372T and Ad-S-D428N in RBD were more potent than that with the wild-type Ad-S in eliciting neutralizing antibodies against the Beta (B.1.351) variant.

**Figure 5.**
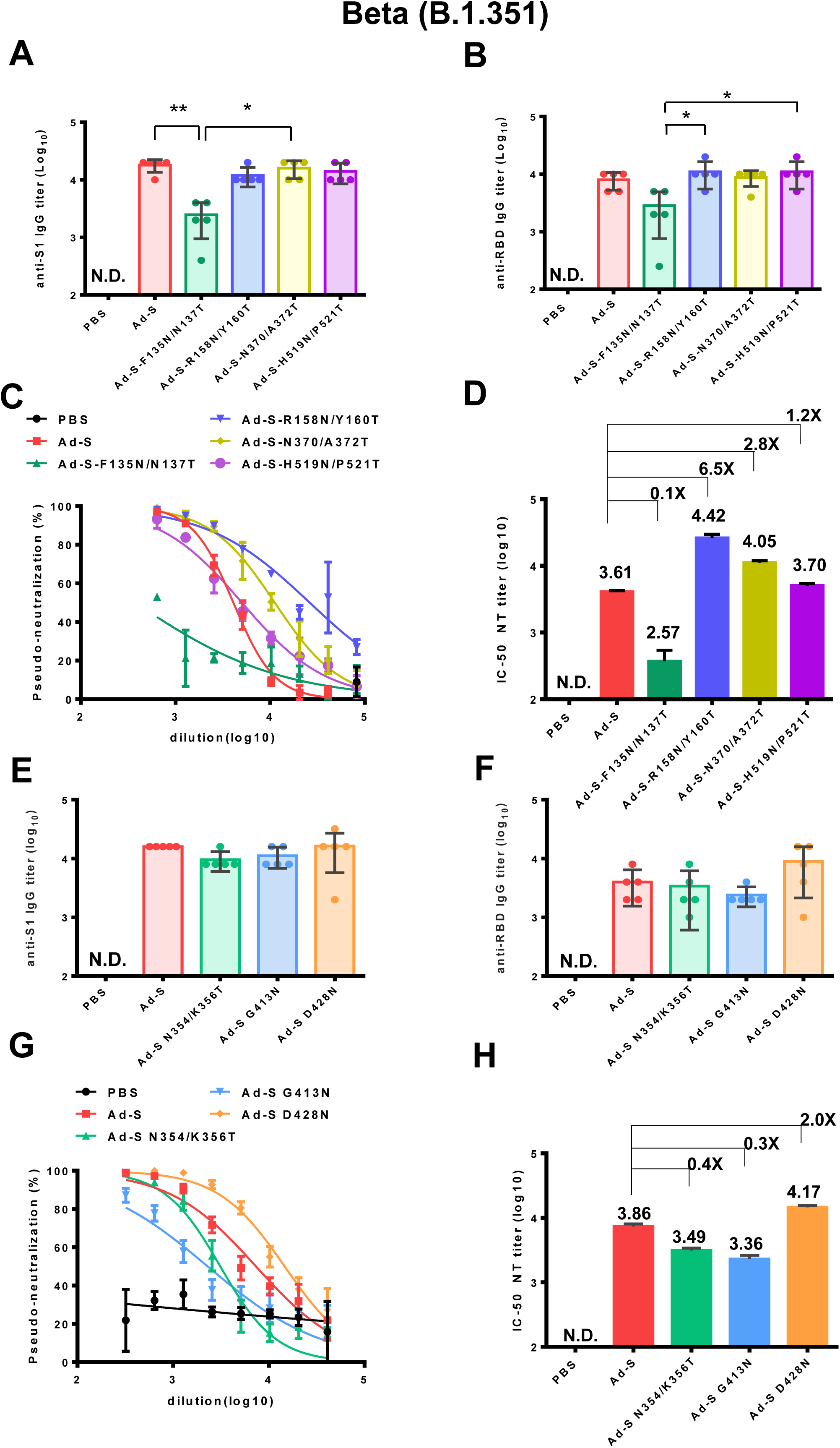
Antibody responses elicited by Ad-S glycan-masking mutants against SARS-CoV-2 Beta (B.1.351) variant. **(A)** antisera for anti-S1 IgG titers from each group of mice (n=5) and tested individually in the first set of immunizations; **(B)** antisera for anti-RBD IgG titers from each group of mice (n=5) and tested individually in the first set of immunization experiments; (**C)** the dose-response pseudo-neutralization curves for the pooled sera of each immunized group of mice (n=5) and measured in triplicate for the first set of immunization experiments; (**D)** IC-50 NT titer for neutralizing antibodies against Beta (B.1.351 variant) for the first set of immunization experiments; **(E)** antisera for anti-S1 IgG titers from each group of mice (n=5) and tested individually in the second set of immunizations; **(F)** antisera for anti-RBD IgG titers from each group of mice (n=5) and tested individually in the second set of immunization experiments; (**G)** the dose-response pseudo-neutralization curves for the pooled sera of each immunized group of mice (n=5) and measured in triplicate for the second set of immunization experiments; (**H)** IC-50 NT titer for neutralizing antibodies against Beta (B.1.351 variant) for the first set of immunization experiments. Fold-changes of the Ad-S glycan-masking mutant IC-50 NT titer against the wild-type Ad-S (Beta, B.1.351 variant) titer are indicated. Statistical tests for multiple comparison of anti-S1 and anti-RBD IgG titers among Ad-s immunization groups were performed using the nonparametric test with Kruskal-Wallis with corrected Dunn’s multiple comparison. The statistical significance is expressed as follows: *p < 0.05 and **p < 0.01. Neutralization curves were fitted based on the equation of nonlinear regression log (inhibitor) vs. normalized response -- variable slope using GraphPad Prism v6.01. The IC50 values of neutralization were obtained from the fitting curves using GraphPad Prism v6.01. Error bars are plotted as standard deviation from the mean value. Not detectable for N.D.

In the case of Delta (B.1.617.2) variant, we found that the anti-S1 IgG titers in the Ad-S-F135N/N137T-immunized group were lower than the wild type Ad-S and Ad-S-R158N/Y160T,-immunized groups (**Fig. 6A**). No significant differences in anti-RBD IgG titers were observed among these Ad-immunized groups in the first set of experiments (**Fig. 6B**). Both the Ad-S-F135N/N137T and Ad-S-R158N/Y160T-immunized groups elicited more potent pseudo-neutralization against the Delta (B.1.167.2) variant, as compared to the Ad-S, Ad-S-N370/A372T, and Ad-S-H519N/P521T-immunized groups, in the first set of immunization experiments (**Fig. 6C**); an approximately 3.7-fold and 4.6-fold increase in the neutralization IC50 titer was found for the glycan-masking Ad-S-F135N/N137T and Ad-S-R158N/Y160T groups, respectively (**Fig. 6D**). In the second set of immunization experiments among the Ad-S, Ad-S-N354/K356T, Ad-S-G413N, and Ad-S-D428N-immunized groups, the pseudo-neutralization curves of the three glycan-masking groups against the Delta (B.167.2) variant were less potent than that of the wild-type Ad-S-immunized group (**Fig. 6G**) with a reduced IC50 titers to 0.46, 0.23, and 0.46 fold, respectively (**Fig. 6H**). Therefore, only the glycan-masking Ad-S-F135N/N137T and Ad-S-R158N/Y160T in NTD elicited more potent neutralizing antibody titers against the Delta (B.1.617.2) variant.

**Figure 6.**
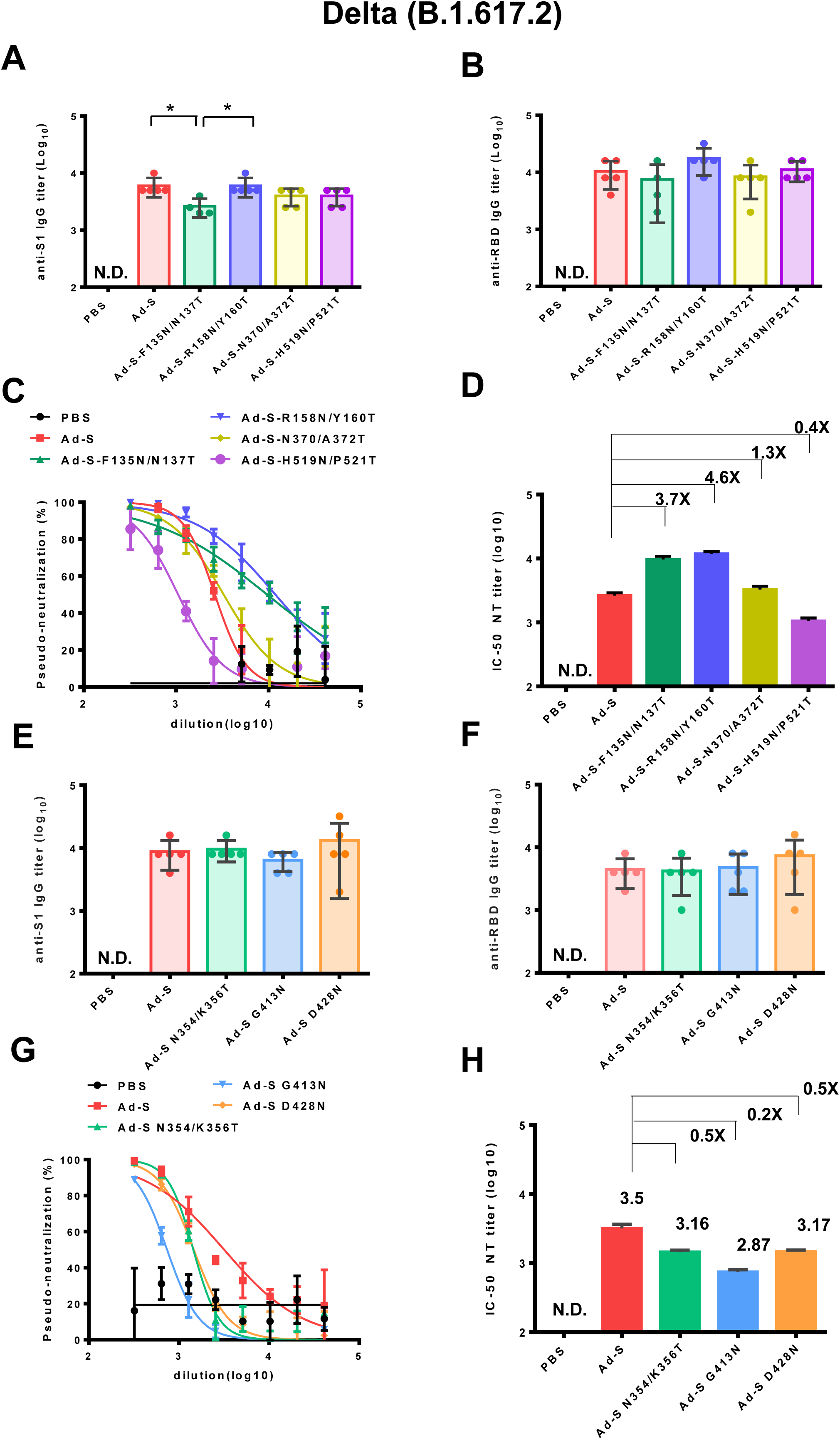
Antibody responses elicited by Ad-S glycan-masking mutants against SARS-CoV-2 Delta (B.1.617.2) variant. **(A)** antisera for anti-S1 IgG titers from each group of mice (n=5) and tested individually in the first set of immunization experiments; **(B)** antisera for anti-RBD IgG titers from each group of mice (n=5) and tested individually in the first set of immunization experiments; (**C)** the dose-response pseudo-neutralization curves for the pooled sera of each immunized group of mice (n=5) and measured in triplicate for the first set of immunization experiments; (**D)** IC50 NT titer for neutralizing antibodies against Delta (B.1.617.2 variant) in the first set of immunization experiments; **(E)** antisera for anti-S1 IgG titers from each group of mice (n=5) and tested individually in the second set of immunization experiments; **(F)** antisera for anti-RBD IgG titers from each group of mice (n=5) and tested individually in the second set of immunization experiments; (**G)** the dose-response pseudo-neutralization curves for the pooled sera of each immunized group of mice (n=5) and measured in triplicate for the second set of immunization experiments; (**H)** IC50 NT titer for neutralizing antibodies against Delta (B.1.617.2 variant) in the second set of immunization experiments. Fold-changes of the Ad-S glycan-masking mutant IC50 NT titer against the wild-type Ad-S (Delta, B.1.617.2 variant) titer are indicated. Statistical tests for multiple comparison of anti-S1 and anti-RBD IgG titers among Ad-S immunization groups were performed using the nonparametric test with Kruskal-Wallis with corrected Dunn’s multiple comparison. The statistical significance is expressed as follows: *p < 0.05. Neutralization curves were fitted based on the equation of nonlinear regression log (inhibitor) vs. normalized response -- variable slope using GraphPad Prism v6.01. The IC50 values of neutralization were obtained from the fitting curves using GraphPad Prism v6.01. Error bars are plotted as standard deviation from the mean value. Not detectable for N.D.

### Comparison of neutralization IC50 titers elicited by glycan-masking Ad-S mutants against the Wuhan-Hu-1 ancestral strain

To compare these results, the neutralizing IC50 titers from the two separate sets of immunization experiments were normalized to the titer elicited by the wild-type Ad-S against the Wuhan-Hu-1 ancestral strain from. In the first set of immunization experiments, the neutralization IC50 titers elicited in the glycan-masking Ad-S-R158N/Y160T-immunized group showed a 2.5-fold increase against the Wuhan-Hu-1 ancestral strain, a 1.8-fold increase against the Alpha (B.1.1.7) variant, a 1.2-fold increase against the Beta (B.1.351) variant, but a 0.6-fold decrease against the Delta (B.1.617.2) variant (**Fig. 7A**). The titer for the Ad-S-R158N/Y160T group against the Delta (B.1.617.2) variant was still higher than the titer of Ad-S against the Delta variant (**Fig. 7A**). In the second set of immunization experiments, the neutralization IC50 titers elicited in the Ad-S-D428N-immunized group showed a 2.7-fold increase against the Wuhan-Hu-1 ancestral strain, a 3.2-fold increase against the Alpha (B.1.1.7) variant, a 2.0-fold increase against the Beta (B.1.351) variant, but a 0.2-fold decrease against the Delta (B.1.617.2) variant (**Fig. 7B**). Therefore, the glycan-masking Ad-S-R158N/Y160T in NTD elicited more potent neutralizing antibodies against the Wuhan-Hu-1 ancestral strain and increased the cross-neutralizing antibody titers against the Alpha (B.1.1.7), Beta (B.1.351), and Delta (B.1.617.2) variants. The glycan-masking Ad-S-D428N in RBD also elicited more potent neutralizing antibodies against the Wuhan-Hu-1 ancestral strain but only increased the cross-neutralizing antibody titers against the Alpha (B.1.1.7) and Beta (B.1.351) variants.

**Figure 7.**
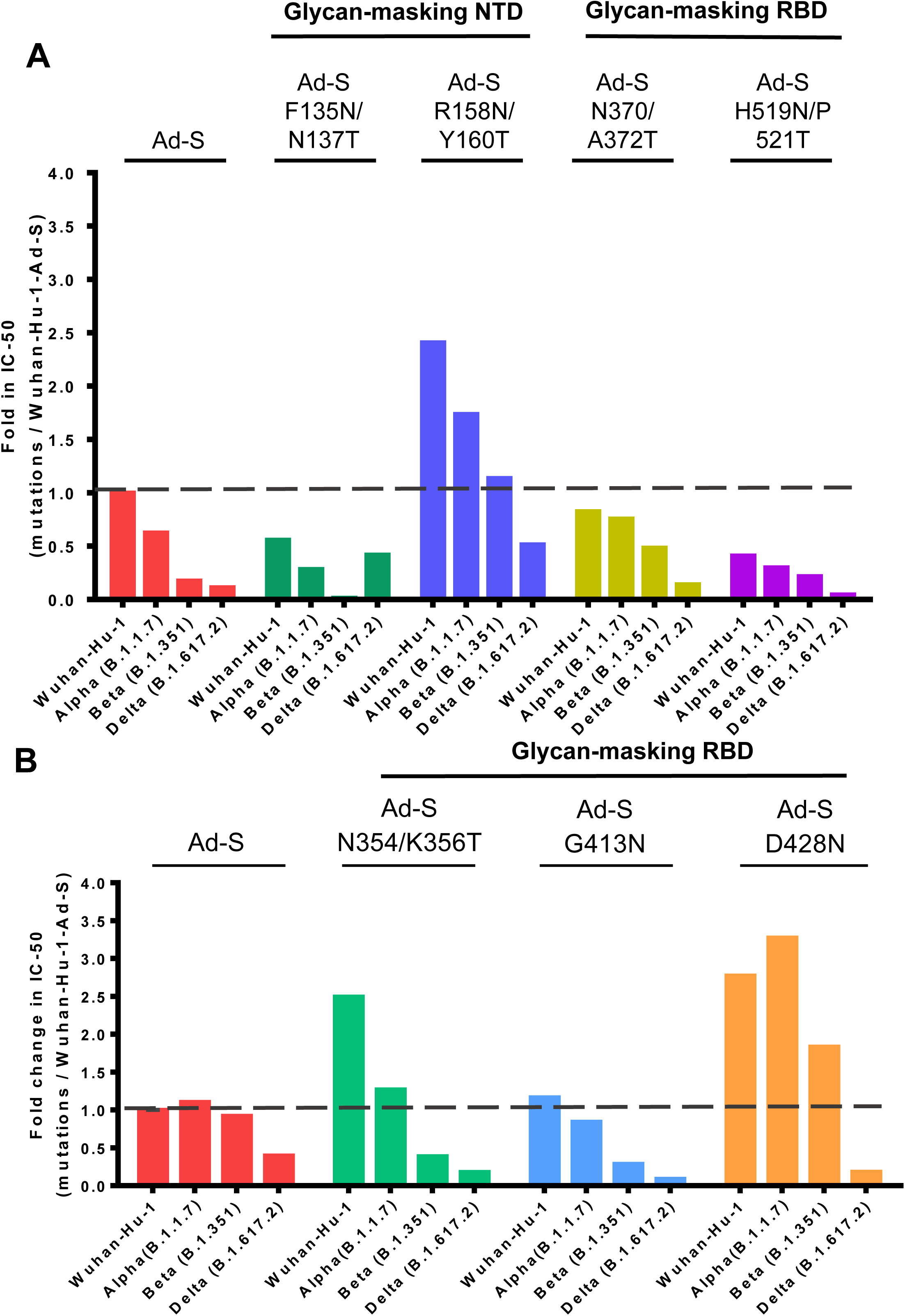
Comparison of neutralization IC50 titers elicited by glycan-masking Ad-S mutants against the Wuhan-Hu-1 ancestral strain. **(A)** Fold-changes in the IC50 NT titers against the Wuhan-Hu-1 ancestral strain in the first set of immunization experiments (5 × 10^7^ pfu per dose) with the glycan-masking Ad-S-F135N/N137T, Ad-S-R158N/Y160T, Ad-S-N370/A372T, and Ad-S-H519N/P521T-immunized groups, as compared to the IC50 titer of the wild-type Ad-S-immunized group; **(B)** Fold-changes in the IC50 NT titers against the Wuhan-Hu-1 ancestral strain in the second set of immunization experiments (1 x10^8^ pfu per dose) with the glycan-masking Ad-S-N354/K356T, Ad-S-G413N, and Ad-S-D428N-immunized groups, as compared to the IC50 titer of the wild-type Ad-S-immunized group.

## Discussion

This study reported the use of glycan-masking antigen design by selective incorporation of an N-linked glycosylation motif on the NTD and RBD in the SARS-CoV-2 S gene encoded by the Ad vector, to develop universal COVID-19 vaccines. It has been recently confirmed that the S protein of SARS-CoV-2 is heavily glycosylated, with approximately 14-16 or 17 N-linked N-glycans out of 22 potential N-glycan sites on each S monomer (30, 32-33). We introduced seven separate N-linked glycosylation sites into the S glycoprotein, S-F135N/N137T, S-R158N/Y160T, Ad-S-N354/K356T, S-N370/A372T, Ad-S-G413N, Ad-S-D428N and S-H519N/P521T, in the NTD and RBD. However, we were unable to demonstrate the addition of a single N-glycan on the target sites for these glycan-masking Ad-S mutants using SDS-PAGE gel without and with PNGase treatment in western blots, to show an increase in the molecular weights of the glycan-masking mutants, as compared to that of the wild-type Ad-S. It is also likely that glycan-masking mutations may also affect the S protein stabilization for cell surface expression, S/S1 cleavage, and surface S expression. Thus, there is a need for further characterization of these glycan-masking mutants, particularly Ad-S-R158N/Y160T and Ad-S-G413N expressed S proteins.

Our results showed that the glycan-masking Ad-S-R158N/Y160T at the N3 loop in the NTD and the glycan-masking Ad-S-N354/K356T and Ad-S-G413N at the C-3 and C-7 loops in the RBD (**Fig. S1**) elicited a potent neutralizing antibody response against the Wuhan-Hu-1 ancestral strain (**Fig. 3**). Selection of these glycan-masking sites in this investigation was based on visual inspection of the 3-D S protein structure (PDB ID: 7C2L) for the exposed loops or the protruding sites of the exposed loops in NTD and RBD of the S1 subunit (**Fig. S1**). The increased IC50 NT titers against the Wuhan-Hu-1 ancestral strain by the glycan-masking Ad-S-R158N/Y160T and Ad-S-G413N-immunized groups correlated with the increased neutralization titers against the Alpha (B.1.1.7) and Beta (B.1.351) variants, but not the neutralization titers against the Delta (B.1.617.2) variant (**Figs. 4-6**). Only the glycan-masking Ad-S-F135N/N137T and Ad-S-R158N/Y160T in NTD were found to increase the neutralization titers against the Delta (B.1.617.2) variant (**Fig. 6D**). Therefore, only the glycan-masking Ad-S-R158N/Y160T in NTD elicited broadly neutralizing antibody titers against Alpha (B.1.1.7), Beta (B.1.351), and Delta (B.1.617.2) variants. It is likely that the refocusing antibodies using the glycan-masking Ad-S-R158N/Y160T antigen may target the NTD neutralizing epitopes as recently reported (34-35). It is also likely that the glycan-masking R158N/Y160T in the NTD interacts spatially with the RBD of another S1 monomer to affect the RBD up and down conformational structures (36-37). The C-type lectins such as L-SIGN and DC-SIGN have been shown to function as attachment receptors by enhancing ACE2-mediated infection, and monoclonal antibodies to NTD or the RBD conserved site at can effectively block lectin-facilitated infection (38). Additionally, the unique N-glycan on N149 of NTD can directly bind to the L-SIGN/DC-SIGN lectins as a non-ACE2 receptor for SARS-CoV-2 virus infection (39). Another report was shown for the N92 glycan on NTD that can enhance the binding to the L-SIGN lectin to interact with the ACE2 receptor to further facilitate SARS-CoV-2 virus entry (40). It is likely that the glycan-masking Ad-S-R158N/Y160T site which is nearby the N3 loop on NTD (**Fig. S1**) can enhance targeting these epitopes to elicit neutralizing antibodies to block the L-SIGN/DC-SIGN receptor binding and/or the interaction between the L-SIGN/DC-SIGN lectin co-receptor with the ACE2 receptor binding for SARS-CoV-2 infection.

Our present findings demonstrated that the glycan-masking Ad-S-R158N/Y160T in NTD resulted in a 2.8-fold, 6.5-fold, and 4.6-fold increase, respectively, in the IC50 titers against the Alpha (B.1.1.7), Beta (B.1.351) and Delta (B.1.617.2) variants, respectively. The glycan-masking Ad-S-D428N in RBD resulted in a 3.0-fold and 2.0-fold increase, respectively, in the IC-50 NT titer against the Alpha (B.1.1.7) and Beta (B.1.351) variants. The glycan-masking Ad-S-R158N/Y160T site is close to the del 156-157 of the Delta (B.1.617.2) variant and the del 143 and 144V mutation of the Alpha (B.1.1.7) in the N3 loop on NTD (**Fig. S1**). The glycan-masking Ad-S-D428N was nearby the K417N mutation of the Beta (B.1.351) variant in the C7 loop on RBD (**Fig. S1**). For the Alpha (B.1.1.7) and Beta (B.1.351) variants, the del 69-70, del 144, and del 242-244 deletions in NTD and the K417N/T, E484K, and N501Y mutations in RBD have been shown to increase ACE2 binding affinity and evade antibody-mediated immunity (7-15). It is likely that selective pressures on the NTD and RBD epitopes of the S1 subunit may ultimately result in immune-evasion variants. Our present findings demonstrated the use of glycan-masking mutations in the neutralization-sensitive NTD and RBD epitopes of S1 subunit can refocus antibody responses to the broadly neutralizing epitope domains to overcome the immune-evasion variants. Therefore, glycan-masking the site-specific NTD and RBD epitopes may help develop universal COVID-19 vaccines against current and future emerging SARS-CoV-2 variants.

## Authors’ contributions

conceptualization WSL, ICC, HCC, SCW; formal analysis WSL, ICC, HCC; funding acquisition YCL, SCW; investigation WSL, ICC, HCC; supervision SCW; writing SCW: all authors provided feedback to the final draft.

## Data Availability Statement

The raw data supporting the conclusions of this article will be made available by the authors, without undue reservation.

## Ethics Statement

All experiments were conducted in accordance with the guidelines of the Laboratory Animal Center of the National Tsing Hua University (NTHU). Animal use protocols were reviewed and approved by the NTHU Institutional Animal Care and Use Committee (approval no. 109047)

## Acknowledgements

We thank the RNAi core facility at Academia Sinica for performing the SARS-CoV-2 S-pseudotyped neutralization assay.

## Supplementary Material

The Supplementary Material for this article can be found on line at :

**Supplementary Figure 1.**
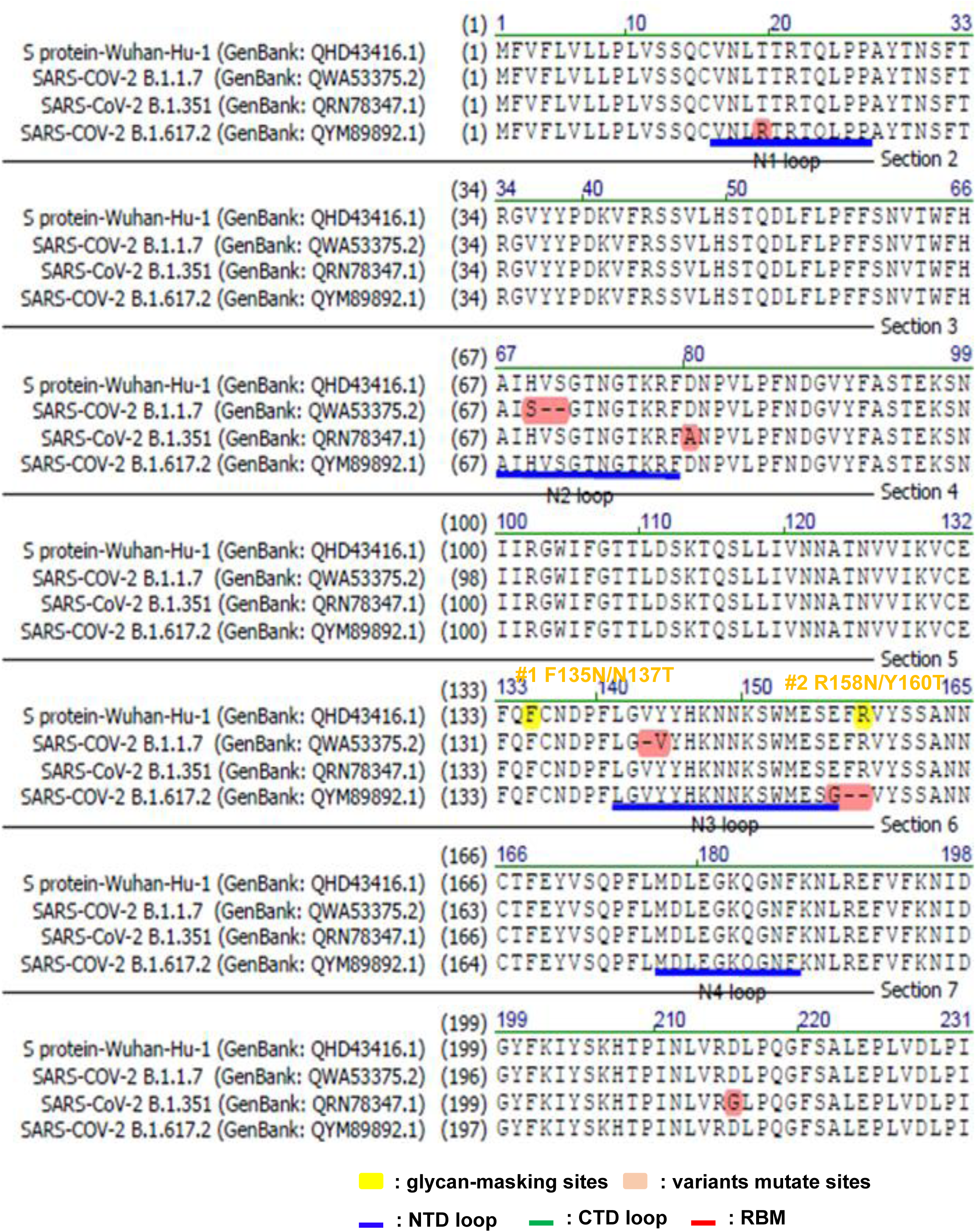
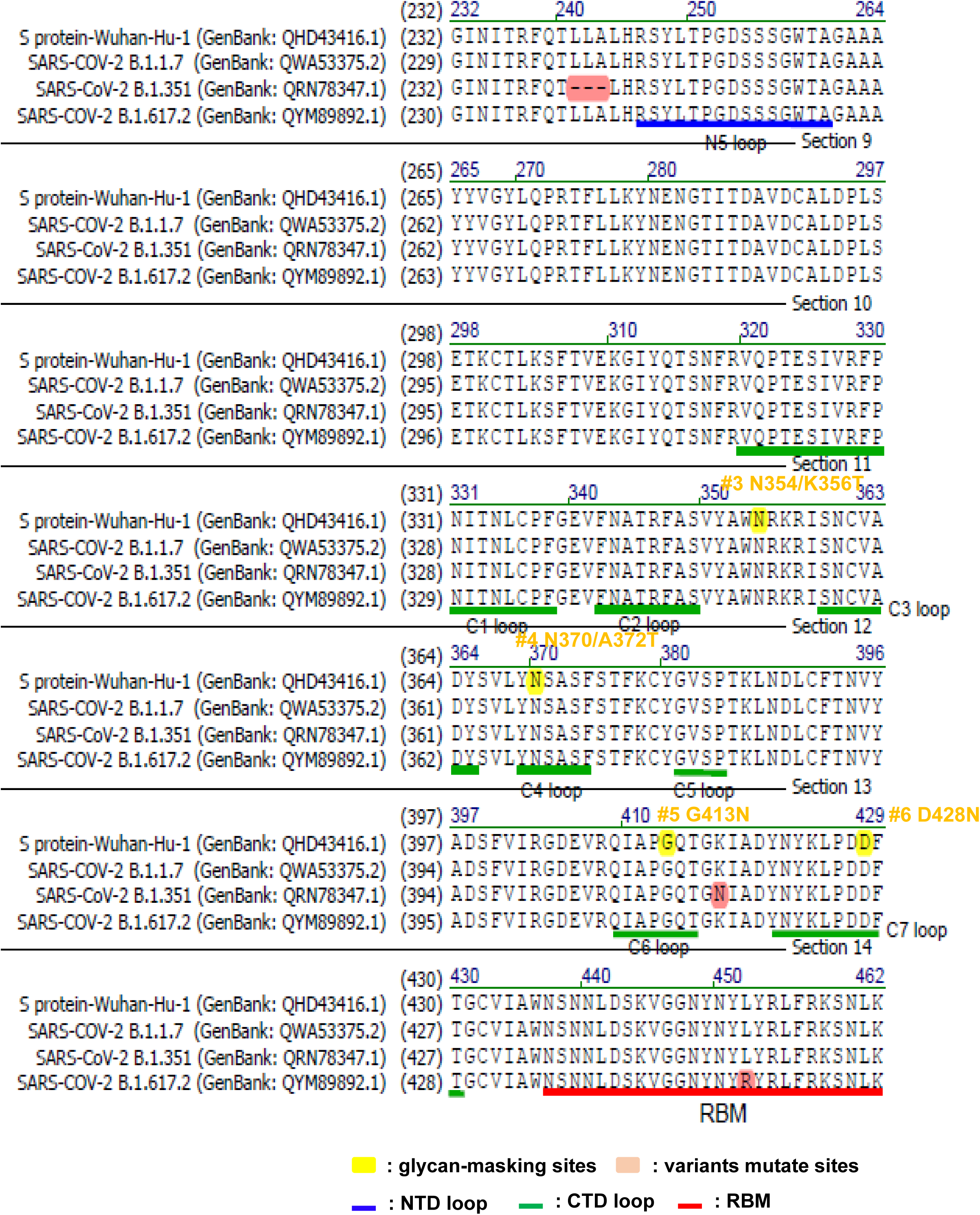
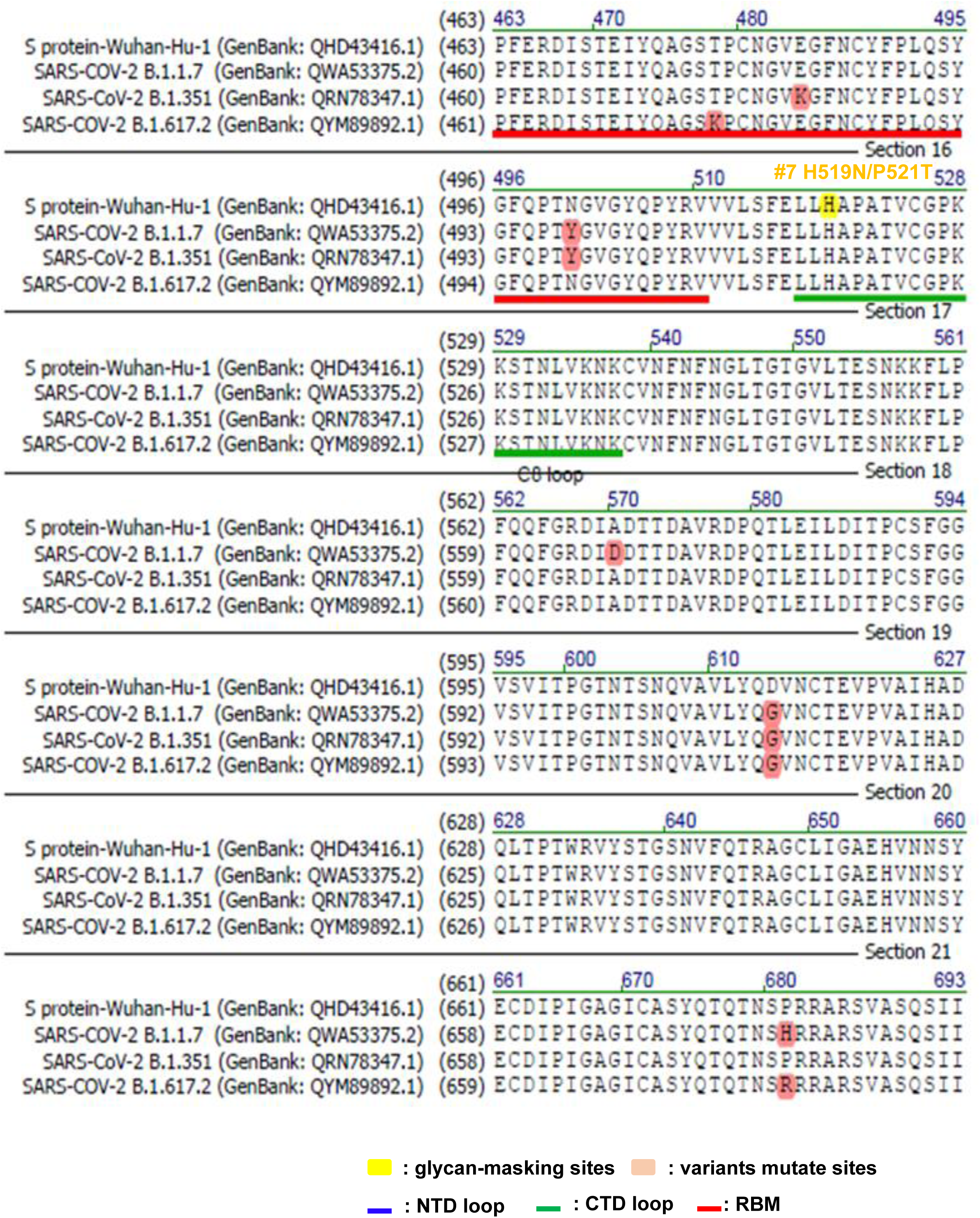
Amino acid sequence alignment of SARS-CoV-2 Wuhan-Hu-1, Alpha (B.1.1.7), Beta (B.1.351), and Delta (B.1.617.2) strains. The NTD loops (blue), the RBD loops (green), and the receptor-binding motif (RBM, red) in the S protein are indicated. The glycan-masking sites (yellow) and variant mutation sites (pink) are marked.

